# Causal relationships between blood lipids and depression phenotypes: A Mendelian randomization analysis

**DOI:** 10.1101/363119

**Authors:** Hon-Cheong So, Carlos Kwan-long Chau, Yu-ying Cheng, Pak C. Sham

## Abstract

**Background:** The etiology of depression remains poorly understood. Changes in blood lipid levels were reported to be associated with depression and suicide, however study findings were mixed.

**Methods:** We performed a two-sample Mendelian randomization (MR) analysis to investigate the causal relationship between blood lipids and depression phenotypes, based on large-scale GWAS summary statistics (*N*=188,577/480,359 for lipid/depression traits respectively). Five depression-related phenotypes were included, namely major depressive disorder (MDD; from PGC), depressive symptoms (DS; from SSGAC), longest duration and number of episodes of low mood, and history of deliberate self-harm (DSH)/suicide (from UK Biobank). MR was conducted with inverse-variance weighted (MR-IVW), Egger and Generalized Summary-data-based MR(GSMR) methods.

**Results:** There was consistent evidence that triglyceride (TG) is causally associated with DS (MR-IVW beta for one-SD increase in TG=0.0346, 95% CI=0.0114-0.0578), supported by MR-IVW and GSMR and multiple r^2^ clumping thresholds. We also observed relatively consistent associations of TG with DSH/suicide (MR-Egger OR= 2.514, CI: 1.579-4.003). There was moderate evidence for positive associations of TG with MDD and the number of episodes of low mood. For HDL-c, we observed moderate evidence for causal associations with DS and MDD. LDL-c and TC did not show robust causal relationships with depression phenotypes, except for weak evidence that LDL-c is inversely related to DSH/suicide. We did not detect significant associations when depression phenotypes were treated as exposures.

**Conclusions:** This study provides evidence to a causal relationship between TG, and to a lesser extent, altered cholesterol levels with depression phenotypes. Further studies on its mechanistic basis and the effects of lipid-lowering therapies are warranted.

## INTRODUCTION

Major depressive disorder (MDD) is one of the most common psychiatric disorders worldwide. It has been ranked as the largest contributor to global disability according to a recent WHO report^1^, and is a major cause underlying suicidal deaths^1^. Nevertheless, the etiology of depression remains largely unclear.

There have been a lot of efforts to search for biological markers associated with and/or causal to depression/suicide, among which serum lipids is one possible etiological factor that is quite widely studied^2-4^. Since dyslipidemia is common and a large number of people are on lipid-lowering therapies^5 6^, the topic is also of high clinical relevance. We shall highlight some epidemiological studies in this area. In an early study, Muldoon et al.^7^ examined the effects on lowering cholesterol concentration on mortality, and revealed a significant rise in deaths due to accidental causes and suicide. Engelberg^8^ proposed that reduced peripheral cholesterol levels may contribute to a decrease in brain serotonin, and that low membrane cholesterol may reduce the number of serotonin receptors, leading to elevated suicidal risks. Subsequently, more clinical studies have been performed; however the results were mixed, with studies showing positive, inverse or non-significant associations^2-4,9^. A few meta-analyses have been performed in this area. Shin et al.^3^ reported that total cholesterol(TC) was inversely associated with levels of depression, especially among drug-naïve patients. They also observed an inverse relationship of low density lipoprotein cholesterol(LDL-c) with depression, but it did not reach statistical significance. Interestingly, they reported a positive association of high density lipoprotein cholesterol(HDL-c) with depression in women. A more recent meta-analysis also reported that lower LDL-c is associated with higher risk of depression^2^. On the other hand, depression may be associated with raised coronary heart disease(CHD) risks^10^ and high LDL-c is a major risk factor for CHD^11^. As for suicidal risks, in a meta-analysis, Wu et al. reported lower serum lipids are generally associated with heightened suicidality^4^. Overall speaking, the findings are mixed and inconsistent, and heterogeneity between studies is large^2-4^.

There are several important limitations in previous studies. One key limitation is that *cause-effect* relationships cannot be reliably determined from previous studies. Many studies are case-control or cross-sectional in nature and the temporal relationship between depression onset and lipid changes cannot be ascertained. In addition, confounding variables may not be completely controlled for in every study, which hinders causal inference. For example, medications, including some antidepressants, may affect the lipid profiles of patients^12^. It is difficult to control for drug effects unless the study only involves drug-naïve cases. In addition, publication bias might be present, and previous meta-analyses did reveal statistical evidence of such bias^2,4^. Although the effects of cholesterol levels on depression were investigated in a number of works, the effects of triglyceride, another major lipid, were less well-studied. Moreover, relatively few studies (except e.g.^13,14^) focused on other depression phenotypes such as duration of depressive episode or symptoms.

In this study, we aim to analyze *causal relationships* between lipid levels and depression-related phenotypes. We will employ Mendelian randomization(MR) for causal inference. MR makes use of genetic variants as “instruments” to represent the exposure of interest, and infers causal relationship between the exposure and the outcome^15^. MR is much less susceptible to confounding bias and reverse causality when compared to observational studies. The principle of MR may be considered similar to a randomized controlled trial(RCT): for example, a group of subjects who have inherited lipid-lowering alleles at a locus (or a set of such alleles at multiple loci) will have lower lipid levels on average, which is analogous to receiving lipid-lowering medication(s) in an RCT^16^. The random allocation of alleles at conception is analogous to random assignment of treatment in an RCT. Another advantage is that MR can be conducted with summary statistics from genome-wide association studies(GWAS), which are commonly of large sample sizes. Here we studied five depression-related phenotypes, including major depressive disorder (MDD), depressive symptoms (DS), longest duration of depressed mood, number of episodes having depressed mood and history of suicide or deliberate self-harm. The aim of studying multiple phenotypes is to gain a comprehensive understanding of the effects of lipids on depression traits and to triangulate the results from different aspects.

## METHODS

For a more detailed description of samples/methods, please also refer to Supplementary Text.

### GWAS study samples

The study samples were primarily of European ancestry. Four lipid traits are studied, including LDL-c, HDL-c, triglyceride(TG) and total cholesterol(TC). GWAS was performed by the Global Lipids Genetics Consortium(*N*=188,577). We downloaded summary statistics of joint GWAS analysis from http://csg.sph.umich.edu/willer/public/lipids2013/. For details please refer to ref^17^.

We included five depression-related phenotypes as follows:

1. Major depression disorder(MDD): We employed the latest PGC GWAS meta-analysis from Wray et al.^18^ *Full* summary statistics are available for a subset of subjects excluding 23andMe participants(59851 cases/113154 controls). This set of summary statistics was used for MR analysis with lipids as exposure. The UK BioBank(UKBB) sub-sample (14260 of 59851 cases) included some cases from self-reporting, while others were defined by clinical assessment/records. We also performed MR with MDD as exposure, for which we used the ‘top 10k SNPs’ from the entire sample(135458 cases/344901 controls). Cases from 23andMe were based on self-reporting.
2. Depressive symptoms(DS): GWAS results were taken from^19^, a meta-analysis from SSGAC that included the MDD-PGC study(*N*=18,759), a case-control sample from the GERA Cohort(*N*=56,368), and a UKBB *general population* sample(*N*=105,739, 58% of total sample). DS were measured by a self-reported questionnaire in UKBB.

GWAS of phenotypes (3) to (5) were based on UKBB. We downloaded GWAS summary statistics from the Neale Lab(https://sites.google.com/broadinstitute.org/ukbbgwasresults/). The analytic approach is given in https://github.com/Nealelab/UK_Biobank_GWAS/tree/master/imputed-v2-gwas and http://www.nealelab.is/blog/2017/9/11/details-and-considerations-of-the-uk-biobank-gwas. For binary outcome(phenotype 5), we converted the regression coefficients and their SE under the linear model to those under a logistic model^20^. Phenotypes 3 and 4 were recorded for those who indicated feeling depressed or down for >1 week only.

(3) Longest period of depressed/low mood: This item was based on self-reporting(*N*=104,190). Inverse-rank normal transformation was used.
(4) Number of episodes with depressed/low mood: It was based on response to the question ‘How many periods have you had when you were feeling depressed or down for at least a whole week?’(*N*=104,190).
(5) History of deliberate self-harm (DSH) or suicide: This item was based on self-reporting. There were 224 positive responses among 381,462 participants(https://biobank.ctsu.ox.ac.uk/crystal/field.cgi?id=20002). The low number of positive cases may be due to under-reporting. The controls are likely mixed with positive cases, and may be considered weakly ‘screened’ or almost ‘unscreened’. This does not render the analysis invalid, and the use of unscreened subjects is common in GWAS^21^. However the power of the study will be improved if reporting bias can be eliminated.

There is no overlap between the GWAS samples for lipids and depression phenotypes, also supported by the non-significant intercepts from genetic correlation analysis(Table S3).

### Mendelian randomization(MR) analysis

Here we performed two-sample MR in which the instrument-exposure and instrument-outcome associations were estimated in different samples. We first performed MR with lipids as exposure and depression phenotypes as outcome, then conducted MR in the reverse direction(for 3 out of 5 phenotypes, as there are insufficient SNPs for the other phenotypes).

We conducted MR with several different methods, including the ‘inverse-variance weighted’(MR-IVW)^22^, Egger regression(MR-Egger)^23^ and Generalized Summary-data-based Mendelian Randomization(GSMR)^24^ approaches. One of the concerns of MR is horizontal pleiotropy, in which the genetic instruments have effects on the outcome other than through effects on the exposure. MR-Egger and GSMR are able to give valid estimates of causal effects in the presence of imbalanced horizontal pleiotropy.

#### MR-IVW and MR-Egger accounting for SNP correlations

The IVW framework is widely used in MR. Here we used an IVW approach that is *able to account for SNP correlations*, described in Burgess et al.^25^. A similar approach may be used for MR-Egger, which allows an intercept term in the weighted regression. Please refer to^23,26^ and Supplementary Text for details. The presence of imbalanced horizontal pleiotropy could be assessed by whether the intercept term is significantly different from zero.

Inclusion of a larger panel of SNPs in partial LD may enable higher variance to be explained, thus improving the power of MR^25^. Including “redundant” SNPs in addition to the causal variant(s) do not improve power but will not invalidate the results^25^. However, including too many variants with high correlations may result in unstable causal estimates^27^.

Here we performed LD-clumping of genetic instruments at multiple *r*^2^ thresholds while accounting for their correlations. Burgess et al.^25^ showed in extensive simulations that if SNP correlations are properly accounted for by the above method, *type I error is properly controlled*. The effect size estimate is also close to the true causal effect. Similarly, for another MR method(GSMR; see below) also employed, simulations have shown *good control of type I error* in the presence of correlated instruments^24^.

Conventionally, MR is performed on (approximately) independent SNPs, and hence many studies and the web-program MR-base employed a LD-clumping threshold of 0.01/0.001 by default. However, MR-base does not have a built-in function to account for LD between SNPs (yet), therefore the program requires input SNPs to be independent. If LD is not adjusted for, it may result in ‘over-precise’ estimates(erroneously low SE)^25^. However, with proper ways to deal with LD, relaxation of the *r*^2^ threshold is legitimate and valid as discussed above.

Only SNPs passing genome-wide significance(*p*<5E-8) were included as instruments. Analysis was performed with the R packages “MendelianRandomization”(ver 0.4.1)^28^ and “TwoSampleMR”(ver 4.25)^29^. If a SNP was not available in the outcome GWAS, we allow using a “proxy SNP” provided *r*^2^>=0.8 with the original SNP. LD was extracted from the 1000 Genomes European samples.

#### GSMR approach

Another analytic framework, GSMR(http://cnsgenomics.com/software/gsmr/), also accounts for horizontal pleiotropy but operates based on excluding ‘outlier’ or heterogeneous genetic instruments that are likely pleiotropic(‘HEIDI-outlier’ method)^24^. GSMR also employed a slightly different formula from MR-IVW by modelling variance of both 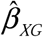 and 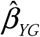. Correlated variants can be accommodated and validity is proven in extensive simulations^24^.

For analysis with <3 genetic instruments, we employed MR-IVW since MR-Egger and GSMR cannot be reliably performed. For single genetic instrument, the Wald ratio approach was used. We also performed the Steiger’s test of directionality^30^ to further ascertain the direction of causal associations.

#### Multivariable MR

In addition to the standard MR which assesses the causal relationship between an outcome and one exposure variable, we also performed multivariable MR(MVMR) to examine the ‘independent’ causal association of each type of lipid(LDL-c/HDL-c/TG) to depression phenotypes, controlling for other lipids. Multivariable MR was performed according to^31,32^. We also performed multivariable MR-Egger(code from^33^) which accounts for imbalanced pleiotropy^33^. The interpretation of univariable and multivariable MR estimates are different. Univariable MR gives the *total* causal effect of an exposure on the outcome, while MVMR gives the ‘independent’ effect of a risk factor adjusting for other modelled exposures. It has been argued the *total* causal effect is more clinically relevant as this is the effect expected from an intervention(e.g. a drug) on the exposure^34^. Another point to note is that methods to adjust for instrument correlations have not been developed for MVMR, therefore the results(especially SEs) at higher correlations should be viewed with caution. For these reasons MVMR is regarded as a secondary analysis here; the main aim is to give a more comprehensive picture on how each exposure exert an independent effect on the outcome *not* mediating through other modelled exposures.

#### LD score regression

We also performed LD score regression(LDSR)^35^ to compute genetic correlations between lipid and depression-related traits. LDSR evaluates the genetic overlap between traits and provides additional support to association between the disorders, although this approach was not designed for assessing causality.

#### Multiple testing control by FDR

Multiple testing was controlled by the Benjamini-Hochberg^36^ false discovery rate(FDR) approach, which controls the expected *proportion* of false positives among the rejected hypotheses. In this study we set a FDR threshold of 0.05 to declare significance, while results with FDR <0.1 are regarded as suggestive associations. FDR control is also valid under positive (regression) dependency of hypothesis tests^37^.

#### ‘Negative control’ analysis

To further ensure the validity of our approach, we performed MR analyses on a ‘negative control’ exposure, assumed to be not causally linked to depression. We chose mean platelet volume^38^ as the exposure and the entire analysis was repeated.

#### General criteria for assessing the reliability of results

We have performed a comprehensive analysis using multiple MR methods and *r*^2^ thresholds. For consistency, we consider an empirical set of criteria to assess the reliability of findings. We consider a finding as having strong evidence or relatively robust if (1)the association is significant/suggestive after multiple testing corrections(FDR<0.1), by at least 2 out of the 3 methods(IVW, Egger, GSMR); (2) the direction of association is consistent across the MR methods that yield significant results; and (3)the result is significant/suggestive after FDR correction(FDR<0.1) across *>=*2 clumping thresholds. For associations that fulfil (1) and (3) but supported by only one MR method, we generally consider them findings of moderate evidence. Other findings that are nominally significant(p< 0.05) at >=2 clumping thresholds are considered as having weak evidence in general. We also evaluate each result case by case. We present the full results of our primary MR analysis as a spreadsheet(Table S1) for easy searching.

We highlight a few points here before proceeding to the full results. As we will observe later, a number of associations were observed at higher r2 thresholds but weaker/absent at more stringent thresholds (*e.g*.0.01/0.001). Relaxing the clumping threshold results in inclusion of more genetic instruments, which usually leads to increase in precision(reduction in SE) of the causal estimates^25^. For example, one may see from Table S1 that for the same MR methodology and exposure-outcome pairs, SE usually decreases as r2 threshold is relaxed. At lower thresholds, the absence of associations could sometimes be due to *less* precise effect estimates(leading to *lower* power for the same effect), and such lack of association per se does *not* preclude the existence of genuine causal relationships. Nevertheless, the actual significance level at different clumping thresholds is dependent on both the effect estimate and its SE(hence p-values may not always be lower with lower thresholds). Secondly, some methods may have better power than others in a given scenario. For example, GSMR was reported to achieve better power than MR-IVW or Egger when the number of instruments is large^24^. If we compare MR-IVW and MR-Egger, the SE of causal estimates is usually higher with MR-Egger^39^. Therefore, even if a finding is genuine, it may *not* be detected by all MR methods. Different MR approaches have their own strengths and limitations, and require different assumptions, which is why we employed multiple approaches here. Please refer to other reviews(e.g.^40,41^) for detailed discussions. Of course, a finding is considered more robust if corroborated by numerous methods.

## RESULTS

### Lipids as exposure and depression-related phenotypes as outcome (univariable MR)

#### LDL-c as exposure

Overall we did not observe highly robust causal associations between LDL-c and depression outcomes(Table 1, S1). The only significant finding that passed FDR correction(FDR<0.05) was LDL-c with DSH/suicide using MR-Egger(*r*^2^=0.15; OR for one-SD *decrease* in LDL-c=1.930, 95% CI:1.234–2.758, *p*=3.02E-4, FDR=7.55E-3). The Egger intercept was significant(intercept=0.0280, p=0.003), hence MR-IVW may be biased. As the finding was not supported by GSMR and at other clumping thresholds, we consider the level of reliability as weak to moderate. Besides, nominally significant(p<0.05) inverse associations of LDL-c with MDD were observed with GSMR at two r2 levels(lowest FDR=0.259) and MR-IVW at one r2 level. However, since the result did not pass FDR correction, we consider this association of at most weak evidence requiring further replications.

**Table 1.**
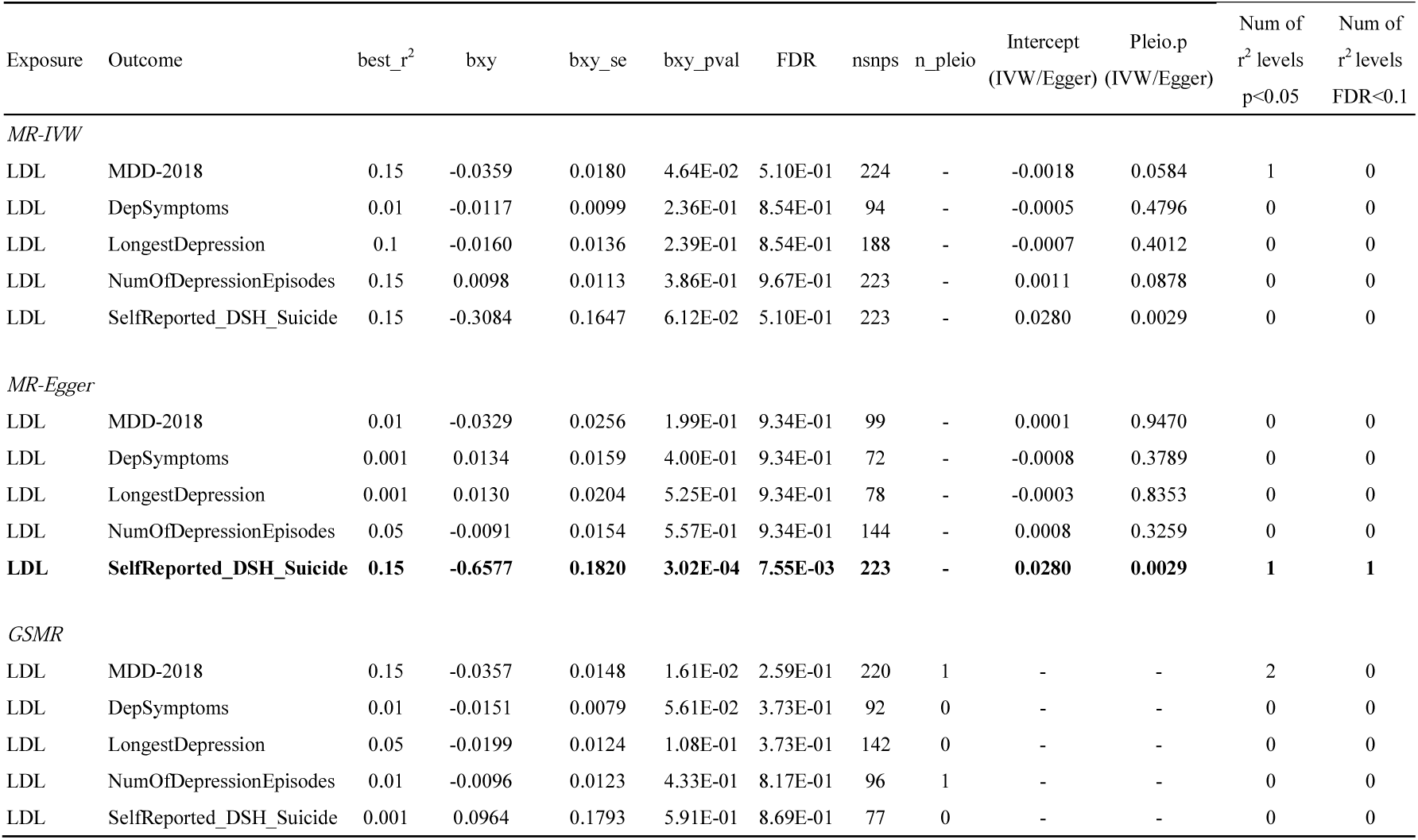
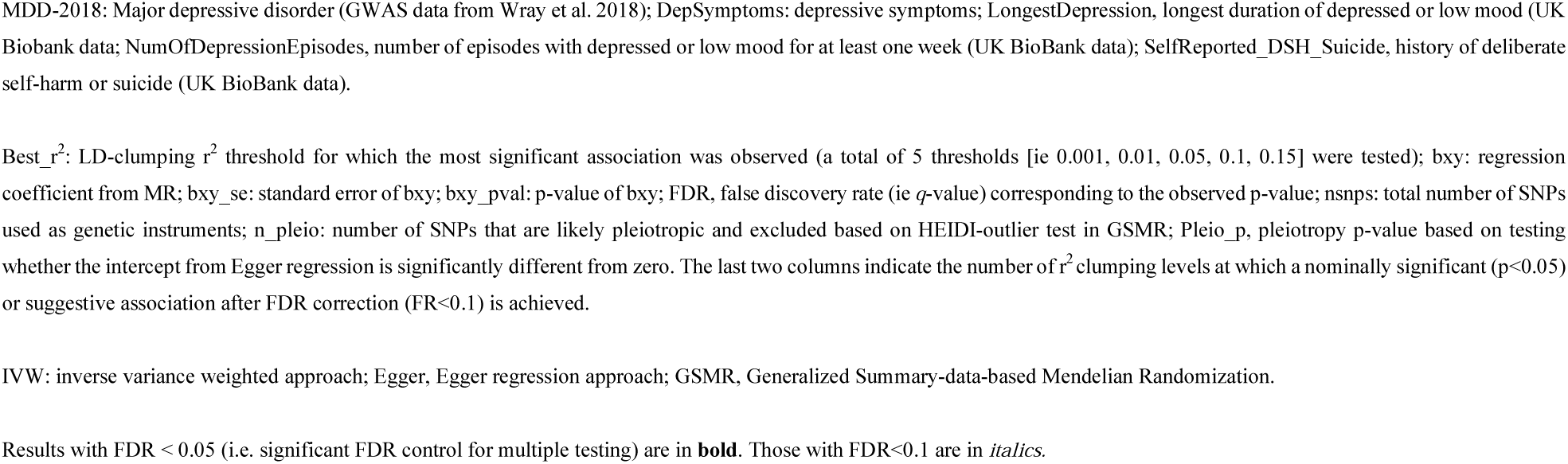
Mendelian randomization (MR) result with **LDL-c** as exposure and depression traits as outcome

| Exposure | Outcome | best_r<sup>2</sup> | bxy | bxy_se | bxy_pval | FDR | nsnps | n_pleio | Intercept<br>(IVW/Egger) | Pleio.p<br>(IVW/Egger) | Num of<br>r<sup>2</sup> levels<br>p<0.05 | Num of<br>r<sup>2</sup> levels<br>FDR<0.1 |
| --- | --- | --- | --- | --- | --- | --- | --- | --- | --- | --- | --- | --- |
| *MR-IVW* |  |  |  |  |  |  |  |  |  |  |  |  |
| LDL | MDD-2018 | 0.15 | -0.0359 | 0.0180 | 4.64E-02 | 5.10E-01 | 224 | - | -0.0018 | 0.0584 | 1 | 0 |
| LDL | DepSymptoms | 0.01 | -0.0117 | 0.0099 | 2.36E-01 | 8.54E-01 | 94 | - | -0.0005 | 0.4796 | 0 | 0 |
| LDL | LongestDepression | 0.1 | -0.0160 | 0.0136 | 2.39E-01 | 8.54E-01 | 188 | - | -0.0007 | 0.4012 | 0 | 0 |
| LDL | NumOfDepressionEpisodes | 0.15 | 0.0098 | 0.0113 | 3.86E-01 | 9.67E-01 | 223 | - | 0.0011 | 0.0878 | 0 | 0 |
| LDL | SelfReported_DSH_Suicide | 0.15 | -0.3084 | 0.1647 | 6.12E-02 | 5.10E-01 | 223 | - | 0.0280 | 0.0029 | 0 | 0 |
| *MR-Egger* |  |  |  |  |  |  |  |  |  |  |  |  |
| LDL | MDD-2018 | 0.01 | -0.0329 | 0.0256 | 1.99E-01 | 9.34E-01 | 99 | - | 0.0001 | 0.9470 | 0 | 0 |
| LDL | DepSymptoms | 0.001 | 0.0134 | 0.0159 | 4.00E-01 | 9.34E-01 | 72 | - | -0.0008 | 0.3789 | 0 | 0 |
| LDL | LongestDepression | 0.001 | 0.0130 | 0.0204 | 5.25E-01 | 9.34E-01 | 78 | - | -0.0003 | 0.8353 | 0 | 0 |
| LDL | NumOfDepressionEpisodes | 0.05 | -0.0091 | 0.0154 | 5.57E-01 | 9.34E-01 | 144 | - | 0.0008 | 0.3259 | 0 | 0 |
| **LDL** | **SelfReported_DSH_Suicide** | **0.15** | **-0.6577** | **0.1820** | **3.02E-04** | **7.55E-03** | **223** | **-** | **0.0280** | **0.0029** | **1** | **1** |
| *GSMR* |  |  |  |  |  |  |  |  |  |  |  |  |
| LDL | MDD-2018 | 0.15 | -0.0357 | 0.0148 | 1.61E-02 | 2.59E-01 | 220 | 1 | - | - | 2 | 0 |
| LDL | DepSymptoms | 0.01 | -0.0151 | 0.0079 | 5.61E-02 | 3.73E-01 | 92 | 0 | - | - | 0 | 0 |
| LDL | LongestDepression | 0.05 | -0.0199 | 0.0124 | 1.08E-01 | 3.73E-01 | 142 | 0 | - | - | 0 | 0 |
| LDL | NumOfDepressionEpisodes | 0.01 | -0.0096 | 0.0123 | 4.33E-01 | 8.17E-01 | 96 | 1 | - | - | 0 | 0 |
| LDL | SelfReported_DSH_Suicide | 0.001 | 0.0964 | 0.1793 | 5.91E-01 | 8.69E-01 | 77 | 0 | - | - | 0 | 0 |

MDD-2018: Major depressive disorder (GWAS data from Wray et al. 2018); DepSymptoms: depressive symptoms; LongestDepression, longest duration of depressed or low mood (UK Biobank data); NumOfDepressionEpisodes, number of episodes with depressed or low mood for at least one week (UK BioBank data); SelfReported\_DSH\_Suicide, history of deliberate self-harm or suicide (UK BioBank data).
Best\_ $r^2$ : LD-clumping $r^2$ threshold for which the most significant association was observed (a total of 5 thresholds [ie 0.001, 0.01, 0.05, 0.1, 0.15] were tested); bxy: regression coefficient from MR; bxy\_se: standard error of bxy; bxy\_pval: p-value of bxy; FDR, false discovery rate (ie $q$ -value) corresponding to the observed p-value; nsnp: total number of SNPs used as genetic instruments; n\_pleio: number of SNPs that are likely pleiotropic and excluded based on HEIDI-outlier test in GSMR; Pleio\_p, pleiotropy p-value based on testing whether the intercept from Egger regression is significantly different from zero. The last two columns indicate the number of $r^2$ clumping levels at which a nominally significant ( $p < 0.05$ ) or suggestive association after FDR correction ( $FR < 0.1$ ) is achieved.
IVW: inverse variance weighted approach; Egger, Egger regression approach; GSMR, Generalized Summary-data-based Mendelian Randomization.
Results with $FDR < 0.05$ (i.e. significant FDR control for multiple testing) are in **bold**. Those with $FDR < 0.1$ are in *italics*.

#### HDL-c as exposure

We observed inverse causal associations between HDL-c and DS with GSMR that passed multiple testing correction(best r2=0.15; beta=0.0244 for one-SD *decrease* in HDL-c, CI:0.0102-0.0387; *p*=7.64E-4; FDR=0.0192) (Table 2, S1). Similar results were observed when the r2 threshold was changed to 0.05 or 0.1, with FDR<0.1(Table S1). On the other hand, using GSMR, we observed a positive association of HDL-c with MDD (best r2=0.001; OR=1.063 for one-SD *increase* in HDL-c, CI:1.018-1.110; *p*=5.6E-3; FDR=0.0467); similar result was observed at r2=0.01. Both of the above associations were supported by an MR method at >=2 thresholds, hence we consider these findings of moderate evidence. Nevertheless, it is slightly surprising that the effect directions are opposite for MDD and DS, which is discussed later.

**Table 2.** MR result with **HDL-c** as exposure and depression traits as outcome

| Exposure | Outcome | best_r<sup>2</sup> | bxy | bxy_se | bxy_pval | FDR | nsnps | n_pleio | Intercept<br>(IVW/Egger) | Pleio.p<br>(IVW/Egger) | Num of r<sup>2</sup><br>levels<br>p<0.05 | Num of r<sup>2</sup><br>levels<br>FDR<0.1 |
| --- | --- | --- | --- | --- | --- | --- | --- | --- | --- | --- | --- | --- |
| *MR-IVW* |  |  |  |  |  |  |  |  |  |  |  |  |
| HDL | MDD-2018 | 0.01 | 0.0431 | 0.0233 | 6.43E-02 | 3.07E-01 | 123 | - | -0.0003 | 0.8462 | 0 | 0 |
| HDL | DepSymptoms | 0.15 | -0.0149 | 0.0080 | 6.33E-02 | 3.07E-01 | 236 | - | -0.0003 | 0.5024 | 0 | 0 |
| HDL | LongestDepression | 0.01 | -0.0149 | 0.0160 | 3.53E-01 | 5.89E-01 | 124 | - | 0.0012 | 0.3556 | 0 | 0 |
| HDL | NumOfDepressionEpisodes | 0.1 | 0.0222 | 0.0129 | 8.59E-02 | 3.07E-01 | 215 | - | 0.0009 | 0.2659 | 0 | 0 |
| HDL | SelfReported_DSH_Suicide | 0.001 | -0.6078 | 0.2310 | 8.50E-03 | 2.13E-01 | 89 | - | -0.0019 | 0.9189 | 1 | 0 |
| *MR-Egger* |  |  |  |  |  |  |  |  |  |  |  |  |
| HDL | MDD-2018 | 0.1 | 0.0364 | 0.0258 | 1.58E-01 | 7.68E-01 | 214 | - | -0.0001 | 0.9141 | 0 | 0 |
| HDL | DepSymptoms | 0.001 | -0.0208 | 0.0179 | 2.46E-01 | 7.68E-01 | 87 | - | 0.0008 | 0.3360 | 0 | 0 |
| HDL | LongestDepression | 0.05 | -0.0428 | 0.0217 | 4.81E-02 | 7.68E-01 | 170 | - | 0.0017 | 0.0806 | 1 | 0 |
| HDL | NumOfDepressionEpisodes | 0.001 | -0.0165 | 0.0295 | 5.78E-01 | 8.63E-01 | 89 | - | 0.0017 | 0.2364 | 0 | 0 |
| HDL | SelfReported_DSH_Suicide | 0.001 | -0.5733 | 0.4010 | 1.53E-01 | 7.68E-01 | 89 | - | -0.0019 | 0.9189 | 0 | 0 |
| *GSMR* |  |  |  |  |  |  |  |  |  |  |  |  |
| **HDL** | **MDD-2018** | **0.001** | **0.0613** | **0.0221** | **5.60E-03** | **4.67E-02** | **86** | **2** | - | - | **2** | **2** |
| **HDL** | **DepSymptoms** | **0.15** | **-0.0244** | **0.0073** | **7.64E-04** | **1.91E-02** | **228** | **7** | - | - | **3** | **3** |
| HDL | LongestDepression | 0.01 | -0.0097 | 0.0151 | 5.23E-01 | 6.89E-01 | 123 | 1 | - | - | 0 | 0 |
| HDL | NumOfDepressionEpisodes | 0.001 | 0.0142 | 0.0164 | 3.87E-01 | 5.55E-01 | 88 | 1 | - | - | 0 | 0 |
| *HDL* | *SelfReported_DSH_Suicide* | *0.001* | *-0.5073* | *0.2206* | *2.15E-02* | *9.37E-02* | *89* | *0* | - | - | *3* | *2* |

Please refer to table 1 for legends.

In addition, we observed weak to moderate evidence of an inverse association of HDL-c with DSH/suicide by GSMR across two clumping thresholds (with FDR<0.1).

#### TC as exposure

Overall no robust causal associations were observed, and no findings passed FDR correction(Table 3, S1). There were nominally significant associations of TC with the number of depressive episodes at two r2 thresholds(MR-IVW), and one nominally significant finding with longest period of depression(GSMR). However, also in view of the absence of associations with other depression phenotypes, we consider the above findings of at most weak evidence.

**Table 3.** MR result with **total cholesterol (TC)** as exposure and depression traits as outcome

| Exposure | Outcome | best_r<sup>2</sup> | bxy | bxy_se | bxy_pval | FDR | nsnps | n_pleio | Intercept<br>(IVW/Egger) | Pleio.p<br>(IVW/Egger) | Num of r<sup>2</sup><br>levels<br>p<0.05 | Num of r<sup>2</sup><br>levels<br>FDR<0.1 |
| --- | --- | --- | --- | --- | --- | --- | --- | --- | --- | --- | --- | --- |
| *MR-IVW* |  |  |  |  |  |  |  |  |  |  |  |  |
| TC | MDD-2018 | 0.15 | -0.0237 | 0.0188 | 2.07E-01 | 6.48E-01 | 267 | - | -0.0008 | 0.4405 | 0 | 0 |
| TC | DepSymptoms | 0.1 | 0.0054 | 0.0079 | 4.95E-01 | 7.96E-01 | 222 | - | -0.0007 | 0.1636 | 0 | 0 |
| TC | LongestDepression | 0.1 | -0.0231 | 0.0122 | 5.89E-02 | 4.91E-01 | 236 | - | -0.0013 | 0.1031 | 0 | 0 |
| TC | NumOfDepressionEpisodes | 0.1 | 0.0212 | 0.0103 | 3.99E-02 | 4.91E-01 | 236 | - | 0.0005 | 0.5195 | 2 | 0 |
| TC | SelfReported_DSH_Suicide | 0.15 | -0.1218 | 0.1482 | 4.11E-01 | 7.96E-01 | 277 | - | -0.0012 | 0.8967 | 0 | 0 |
| *MR-Egger* |  |  |  |  |  |  |  |  |  |  |  |  |
| TC | MDD-2018 | 0.15 | -0.0095 | 0.0236 | 6.88E-01 | 9.92E-01 | 267 | - | -0.0008 | 0.4405 | 0 | 0 |
| TC | DepSymptoms | 0.1 | 0.0190 | 0.0117 | 1.05E-01 | 9.92E-01 | 222 | - | -0.0007 | 0.1636 | 0 | 0 |
| TC | LongestDepression | 0.05 | -0.0153 | 0.0191 | 4.22E-01 | 9.92E-01 | 174 | - | -0.0002 | 0.8651 | 0 | 0 |
| TC | NumOfDepressionEpisodes | 0.1 | 0.0132 | 0.0154 | 3.92E-01 | 9.92E-01 | 236 | - | 0.0005 | 0.5195 | 0 | 0 |
| TC | SelfReported_DSH_Suicide | 0.1 | -0.2706 | 0.2163 | 2.11E-01 | 9.92E-01 | 236 | - | 0.0086 | 0.3893 | 0 | 0 |
| *GSMR* |  |  |  |  |  |  |  |  |  |  |  |  |
| TC | MDD-2018 | 0.1 | -0.0161 | 0.0162 | 3.20E-01 | 8.54E-01 | 218 | 4 | - | - | 0 | 0 |
| TC | DepSymptoms | 0.15 | -0.0076 | 0.0071 | 2.87E-01 | 8.54E-01 | 247 | 6 | - | - | 0 | 0 |
| TC | LongestDepression | 0.1 | -0.0249 | 0.0123 | 4.21E-02 | 5.76E-01 | 229 | 1 | - | - | 1 | 0 |
| TC | NumOfDepressionEpisodes | 0.001 | -0.0131 | 0.0138 | 3.41E-01 | 8.54E-01 | 84 | 1 | - | - | 0 | 0 |
| TC | SelfReported_DSH_Suicide | 0.01 | -0.1744 | 0.1669 | 2.96E-01 | 8.54E-01 | 115 | 2 | - | - | 0 | 0 |

Please refer to table 1 for legends.

#### TG as exposure

We observed a relatively robust causal association between TG and DS (best r2=0.05 for MR-IVW; beta for one-SD increase in TG=0.0346, CI:0.0114-0.0578; *p*=3.48E-3, FDR=0.0483) [Table 4, S1]. The finding was supported by GSMR, with similar effect estimates (best r2=0.1, beta=0.0324, CI:0.0127-0.0519; *p*=1.19E-3, FDR=0.0231). Of note, the association was supported by findings having FDR<0.1 at multiple r2 thresholds. Besides, LDSR showed significant genetic correlation between TC and DS(r_g_=0.138, p=0.004; FDR=0.04). Taken together, there is strong evidence that TG is causally related to higher DS.

**Table 4.** MR result with **triglycerides (TG)** as exposure and depression traits as outcome

| Exposure | Outcome | best_r<sup>2</sup> | bxy | bxy_se | bxy_pval | FDR | nsnps | n_pleio | Intercept<br>(IVW/Egger) | Pleio.p<br>(IVW/Egger) | Num of<br>r<sup>2</sup> levels<br>p<0.05 | Num of<br>r<sup>2</sup> levels<br>FDR<0.1 |
| --- | --- | --- | --- | --- | --- | --- | --- | --- | --- | --- | --- | --- |
| *MR-IVW* |  |  |  |  |  |  |  |  |  |  |  |  |
| *TG* | *MDD-2018* | *0.01* | *0.0516* | *0.0240* | *3.12E-02* | *8.68E-02* | *70* | *-* | *0.0022* | *0.2300* | *2* | *1* |
| **TG** | **DepSymptoms** | **0.05** | **0.0346** | **0.0118** | **3.48E-03** | **4.83E-02** | **98** | **-** | **0.0012** | **0.0850** | **3** | **2** |
| TG | LongestDepression | 0.1 | 0.0177 | 0.0179 | 3.23E-01 | 3.84E-01 | 139 | - | 0.0010 | 0.3307 | 0 | 0 |
| *TG* | *NumOfDepressionEpisodes* | *0.1* | *0.0421* | *0.0183* | *2.11E-02* | *8.57E-02* | *139* | *-* | *0.0001* | *0.9006* | *2* | *2* |
| **TG** | **SelfReported_DSH_Suicide** | **0.15** | **0.5600** | **0.1938** | **3.86E-03** | **4.83E-02** | **178** | **-** | **-0.0240** | **0.0198** | **4** | **4** |
| *MR-Egger* |  |  |  |  |  |  |  |  |  |  |  |  |
| TG | MDD-2018 | 0.1 | 0.0360 | 0.0298 | 2.27E-01 | 6.44E-01 | 125 | - | 0.0008 | 0.5565 | 0 | 0 |
| TG | DepSymptoms | 0.05 | 0.0143 | 0.0158 | 3.65E-01 | 8.55E-01 | 98 | - | 0.0012 | 0.0850 | 0 | 0 |
| TG | LongestDepression | 0.001 | -0.0649 | 0.0362 | 7.35E-02 | 2.62E-01 | 54 | - | 0.0038 | 0.0325 | 0 | 0 |
| TG | NumOfDepressionEpisodes | 0.1 | 0.0402 | 0.0223 | 7.18E-02 | 2.62E-01 | 139 | - | 0.0001 | 0.9006 | 0 | 0 |
| **TG** | **SelfReported_DSH_Suicide** | **0.15** | **0.9221** | **0.2373** | **1.02E-04** | **2.55E-03** | **178** | **-** | **-0.0240** | **0.0198** | **4** | **4** |
| *GSMR* |  |  |  |  |  |  |  |  |  |  |  |  |
| TG | MDD-2018 | 0.1 | 0.0362 | 0.0220 | 1.00E-01 | 1.92E-01 | 122 | 2 | - | - | 0 | 0 |
| **TG** | **DepSymptoms** | **0.1** | **0.0324** | **0.0100** | **1.19E-03** | **2.31E-02** | **119** | **1** | **-** | **-** | **4** | **4** |
| TG | LongestDepression | 0.001 | -0.0151 | 0.0209 | 4.68E-01 | 5.86E-01 | 54 | 0 | - | - | 0 | 0 |
| TG | NumOfDepressionEpisodes | 0.05 | 0.0340 | 0.0164 | 3.79E-02 | 1.16E-01 | 112 | 0 | - | - | 2 | 0 |
| TG | SelfReported_DSH_Suicide | 0.05 | 0.4694 | 0.2170 | 3.05E-02 | 1.16E-01 | 112 | 0 | - | - | 3 | 0 |

Please refer to table 1 for legends.

We found another relatively consistent causal association between TG and DSH/suicide(best r2=0.15; MR-Egger: OR for one-SD increase in TG=2.514, CI: 1.579-4.003; p=1.02E-4; FDR=2.55E-3). The finding was supported by both MR-Egger and MR-IVW at four r2 thresholds(r2=0.01/0.05/0.1/0.15), with FDR<0.1. We note that at r2=0.1 and 0.15, the pleiotropy p-values were significant, indicating imbalanced pleiotropy and MR-IVW could be biased. However, MR-Egger supported significant causal associations at these thresholds. In addition, GSMR detected nominally significant associations at three r2 thresholds, with a relatively low FDR(0.116).

As for other phenotypes, nominally significant associations between TG and MDD were observed at two thresholds by MR-IVW, with FDR less than or close to 0.1(FDR=0.0868 and 0.102 at r2=0.01 and 0.1 respectively). Interestingly, LDSR revealed a highly significant genetic correlation between TG and MDD(rg= 0.129, p = 0.0002, FDR=0.004). The TG->MDD association was supported by one MR approach, and coupled with a significant genetic correlation, we consider the evidence as moderate. For number of depressive episodes, we observed associations at r2=0.1 and 0.15 with MR-IVW, achieving FDR<0.1. It was partially supported by GSMR at two clumping thresholds with FDR close to 0.1. We also consider this finding of moderate evidence.

#### Multivariable MR

MVMR is considered a secondary analysis here. Since there are no rigorous methods to account for SNP correlations in MVMR, we mainly discuss the results at a relatively stringent r2 threshold(0.01), but results at higher thresholds are also presented for reference(Table S2). The results may not be directly comparable to univariable MR where we employed more relaxed thresholds with higher precision in effect estimates(hence potentially better power). At r2=0.01, several results were suggestive at FDR<0.1, including TG->DS, and LDL-c->DS and LDL-c->MDD(all directionally consistent with univariable MR; Egger intercept non-significant). The former(TG->DS) provides support that TG may have a direct causal effect on DS, not entirely mediated via other lipids. The latter associations provide some additional evidence for a causal link between LDL-c and depression, which was weakly supported by univariable MR. Other negative results could be due to lower power with fewer instruments at a lower r2 threshold, and are harder to interpret. However, we note that the associations with DSH/suicide were largely non-significant in MVMR even at higher r2, suggesting that the total causal effect by each lipid type (TG/HDL-c/LDL-c) on DSH/suicide is partially mediated via other lipids.

#### LD score regression

LDSR revealed significant genetic correlations for TG with MDD and DS(Table S3), both of which passed multiple testing correction. Nominally significant genetic correlation was observed for HDL-c and DS. The above findings were all directionally consistent with univariable MR.

#### Steiger Test of Directionality

Table S4 shows the variance explained in the exposure and outcome by the instrument SNPs. The variance explained in the exposure is clearly larger, and the test returns p-values of zero, indicating the causal direction is from lipid traits to depression phenotypes.

#### Results from negative control experiment

As expected, we do not observe MPV being causally associated with depression phenotypes(Table S5). No results passed multiple testing corrections.

### Depression phenotypes as exposure and lipid traits as outcome

To assess whether depression or related phenotypes cause changes in lipid levels, we performed MR analysis in the reversed direction. For depression phenotypes 3 and 4(longest duration and number of episodes of low/depressed mood), there are no genome-wide significant SNPs which also matched to the set of SNPs included in the lipid GWAS; we therefore excluded them from this analysis.

#### MDD as exposure

No results reached significance at FDR<0.05(Table 5, S6). MDD was positively associated with raised TG and reduced LDL-c at nominal significance at several r2 thresholds, however these associations did not survive FDR correction.

**Table 5.** MR result with **depression traits as exposure** and lipids as outcome

| Exposure | Outcome | best_r<sup>2</sup> | bxy | bxy_se | bxy_pval | FDR | nsnps | n_pleio | Intercept<br>(IVW/Egger) | Pleio.p<br>(IVW/Egger) | Num of r2<br>levels<br>p<0.05 | Num of r2<br>levels<br>FDR<0.1 |
| --- | --- | --- | --- | --- | --- | --- | --- | --- | --- | --- | --- | --- |
| *MR-IVW* |  |  |  |  |  |  |  |  |  |  |  |  |
| MDD-2018 | HDL | 0.001 | -0.0233 | 0.0355 | 5.11E-01 | 9.96E-01 | 32 | - | -0.0100 | 0.1493 | 0 | 0 |
| MDD-2018 | LDL | 0.01 | 0.0108 | 0.0365 | 7.66E-01 | 9.96E-01 | 36 | - | 0.0127 | 0.0636 | 0 | 0 |
| MDD-2018 | TC | 0.001 | -0.0121 | 0.0410 | 7.68E-01 | 9.96E-01 | 32 | - | 0.0109 | 0.1528 | 0 | 0 |
| MDD-2018 | TG | 0.001 | 0.0784 | 0.0381 | 3.96E-02 | 3.93E-01 | 32 | - | 0.0083 | 0.2436 | 1 | 0 |
| DepSymptoms | HDL | - | -0.1437 | 0.1538 | 3.50E-01 | 9.33E-01 | 2 | - | - | - | 0 | 0 |
| DepSymptoms | LDL | - | -0.0820 | 0.1647 | 6.19E-01 | 9.80E-01 | 2 | - | - | - | 0 | 0 |
| DepSymptoms | TC | - | 0.0289 | 0.1610 | 8.58E-01 | 9.80E-01 | 2 | - | - | - | 0 | 0 |
| DepSymptoms | TG | - | 0.1592 | 0.1464 | 2.77E-01 | 9.33E-01 | 2 | - | - | - | 0 | 0 |
| SelfReported_DSH_Suicide | HDL | - | -0.022 | 0.0277 | 4.28E-01 | 7.97E-01 | 1 | - | - | - | 0 | 0 |
| SelfReported_DSH_Suicide | LDL | - | 0.0158 | 0.0284 | 5.79E-01 | 7.97E-01 | 1 | - | - | - | 0 | 0 |
| SelfReported_DSH_Suicide | TC | - | -0.0072 | 0.0281 | 7.97E-01 | 7.97E-01 | 1 | - | - | - | 0 | 0 |
| SelfReported_DSH_Suicide | TG | - | -0.0416 | 0.0237 | 7.88E-02 | 7.97E-01 | 1 | - | - | - | 0 | 0 |
| *MR-Egger* |  |  |  |  |  |  |  |  |  |  |  |  |
| MDD-2018 | HDL | 0.001 | 0.2969 | 0.2243 | 1.86E-01 | 3.86E-01 | 32 | - | -0.0100 | 0.1493 | 0 | 0 |
| MDD-2018 | LDL | 0.001 | -0.5795 | 0.2413 | 1.63E-02 | 2.19E-01 | 32 | - | 0.0181 | 0.0153 | 4 | 0 |
| MDD-2018 | TC | 0.001 | -0.3628 | 0.2478 | 1.43E-01 | 3.86E-01 | 32 | - | 0.0109 | 0.1528 | 0 | 0 |
| MDD-2018 | TG | 0.05 | -0.2024 | 0.1923 | 2.93E-01 | 3.90E-01 | 38 | - | 0.0082 | 0.1745 | 0 | 0 |

| GSMR |  |  |  |  |  |  |  |  |  |  |  |  |
| --- | --- | --- | --- | --- | --- | --- | --- | --- | --- | --- | --- | --- |
| MDD-2018 | HDL | 0.001 | -0.0279 | 0.0296 | 3.45E-01 | 9.74E-01 | 32 | 0 | - | - | 0 | 0 |
| MDD-2018 | LDL | 0.01 | 0.0075 | 0.0302 | 8.04E-01 | 9.74E-01 | 36 | 0 | - | - | 0 | 0 |
| MDD-2018 | TC | 0.001 | -0.0140 | 0.0314 | 6.55E-01 | 9.74E-01 | 32 | 0 | - | - | 0 | 0 |
| MDD-2018 | TG | 0.001 | 0.0690 | 0.0290 | 1.74E-02 | 2.33E-01 | 32 | 0 | - | - | 2 | 0 |

Please refer to table 1 for legends. GSMR and MR-Egger are not applicable for DepSymptoms or DSH/suicide as there are too few SNPs. Results are the same at different r<sup>2</sup> thresholds for DepSymptoms and DSH/suicide, so no best r<sup>2</sup> is presented.

#### Depressive symptoms and DSH/suicide as exposure

We did not find any results reaching significance(Table 5). We only have 2 and 1 SNP(s) respectively as instruments for MR analysis of DS and DSH/suicide, hence tests for pleiotropy cannot be carried out.

## DISCUSSIONS

In this study, we employed MR to reveal causal relationships between lipid levels and depression-related phenotypes. For TG, we observed strong evidence for a causal link with increased DS and relatively consistent associations with higher risks of DSH/suicide. There was moderate evidence for positive associations of TG with MDD and number of episodes of low mood.

As for previous studies on TG and depression, a previous meta-analysis^4^ showed that suicidal psychiatric *patients* had lower TG than non-suicidal *patients*, but such difference was not significant when comparing suicidal patients with healthy controls. However, there was significant heterogeneity and evidence of publication bias; also as many studies are cross-sectional or case-control in nature, causality cannot be accurately determined. Our analysis is *not* focused on suicidal risks *within* patients, so is not contradictory with findings from^4^. Some studies reported raised TG in depression(e.g.^42-45^), but meta-analysis on the topic is lacking. Of note, a prospective study in Finnish young adults showed that steeply rising TG levels throughout childhood and adulthood was associated with increased DS in adulthood^43^, consistent with the present finding of a causal role of TG in the development of DS.

For HDL-c, the findings appeared more complex. We observed moderate evidence that raised HDL-c is causally associated with *lower* DS by GSMR. Interestingly, GSMR also revealed a tentative positive association with MDD. Both findings were significant at FDR<0.05. This apparent discordance in causal direction is interesting, and we hypothesize some possible explanations here. Firstly, it is possible that HDL-c may affect subjects at different ‘positions’ of the depression spectrum differently. To recap, the DS GWAS is largely driven by a general population sample (∼58% of entire sample), while patients in the MDD sample were mainly clinically ascertained. A risk factor may affect people with severe DS or clinical MDD differently from those at the lower/middle range of DS(who are *not* clinically depressed). As a related example, Chang et al. showed that the effect of polygenic scores were significantly different at different quantiles of depressive symptom scores^46^. Another possibility for the discrepant direction may be heterogeneity in study samples(detailed later).

A number of epidemiological studies have investigated the link between cholesterol levels and depression. As already mentioned, a recent meta-analysis^3^ reported higher HDL-c was associated with higher MDD risks. However, meta-analyses have also reported inverse associations of LDL-c^2^ and TC^3^ with depression, for which we did not observe robust evidence for causal relationships here. For LDL-c and HDL-c, we observed weak-to-moderate evidence for a causal link with reduced DSH/suicide risk. The finding requires further replications especially due to the small number of cases, but is concordant with meta-analysis^4^ that both types of cholesterol are *lower* in suicidal patients compared to healthy controls.

We also studied causal relationship in the reversed direction, but did not find evidence that depression and related traits cause changes to lipid levels. The number of instrument SNPs included is small especially for DS and DSH/suicide, which may lead to low power in detecting causal relationships. Further analysis in larger samples is warranted.

### Limitations

There are several limitations to the current study. Firstly, there is likely heterogeneity within each study sample. For example, within the sample of MDD patients, they might be substantial differences with respect to disease etiology, symptoms, course of illness etc. Here we have not studied how the causal relationship with lipids may differ with different depression subtypes or symptoms. In a similar vein, we have not studied how the relationship may be affected by patients’ clinical background. For instance, age, sex, baseline metabolic profile, family history, past medical and drug history etc. may all affect the relationship between lipid levels and depression. Future studies may include in-depth analysis on stratified samples(e.g. male/female-only; young/elderly subjects only). Another related limitation is that different covariates may be included in different GWAS studies. However, most have adjusted for population stratification, which is the major confounder in genetic studies.

Yet another limitation is that for studies that included the UKBB, depression traits are self-reported instead of being assessed by health-care professionals by clinical criteria. There may be heterogeneity in the definition of ‘feeling depression/down’ for different people. Of note, phenotypes 3-5 are fully based on self-reporting from participants, while DS and MDD samples are partially based on self-reports. Under-reporting may also be present(especially for DSH/suicide), which limits the effective sample size. Particularly findings concerning DSH/suicide are based on relatively small number of cases, and require further replications to confirm. Despite the above limitations, detailed clinical assessment is very costly and some subjectivity is still inevitable; the current approach allows larger samples to be studied.

In addition, due to the high proportion of subjects on lipid-lowering therapy, genetically predicted lipid levels(especially those at the higher end) may be less correlated with the measured levels than expected. Another potential limitation is collider or selection bias for UKBB sample. The sample appears to be enriched for people who are healthier and better educated^47^; this selection may be associated with both better lipid profiles and lower DS. Such selection bias can lead to bias in MR; but the bias is usually small if the selection effects are weak or moderate^48^. Besides, censoring may be present, in that some people could develop MDD later in life after recruitment.

With regards to the MR approach used in this study, it is also not without limitations. As we employed genetic instruments to model the risk factor, the analysis reflects effects of a *chronic* exposure of (genetically) lowered lipid levels to the outcomes. The effects of shorter-term exposures, such as taking a LDL-lowering drug for 1 year, cannot be inferred with full confidence from MR alone. Also, non-linear relationships between the exposure and outcome are not captured with the present method.

As detailed above, findings from epidemiological studies were mixed and did not completely agree with our findings. Heterogeneity in samples and study design may partially explain this; also, some shared(but non-causal) genetic or environmental risk factors may be present for both depression and dyslipidemia.

### Clinical implications

We highlight some areas of potential clinical relevance here, but we caution that due to various limitations, one should not over-interpret the current findings.

Some may consider that cholesterol or TG levels may serve as predictive biomarkers for depression/suicide. However, MR reflects the effects of long-term exposure and the time-dependent implications are unclear. The effect sizes reported in this study are also relatively modest. If the current findings are replicated, however, blood lipids may be integrated with other biomarkers and clinical factors to construct better prediction models for depression/suicide.

Another potential clinical implication is on whether lipid-lowering therapies may or may not alter depression /suicidal risks. This is a controversial topic and no consistent conclusions have been reached despite numerous studies^49-51,52^. Based on this study, we are unable to conclude whether lipid-lowering therapies result in altered risks of depression/suicide, owing to several limitations. Firstly, MR models the effects of long-term exposure; effects of shorter exposure at later stages of life may not be reliably determined. In addition, different lipid-lowering drugs act in different pathways, and the effects on depression could differ. Also, many lipid-lowering drugs have effects on more than one lipid type, for example statins lower LDL-c but also reduce TG^53^ and raise HDL-c^51^. The combined effect is difficult to judge. Finally, the effects and side-effects of lipid-lowering drugs may differ from patient to patient, and careful judgement of cardiovascular benefit versus other side-effects is important.

From a therapeutic point of view, it will be interesting to study whether TG-lowering therapies may ameliorate DS in patients with comorbid hypertriglyceridemia and depression; however this study cannot provide confirmatory evidence. Of note, a small-scale clinical trial reported that treatment of severe hypertriglyceridemia resulted in improvement of DS^54^.

In conclusion, through an MR analysis with large sample sizes, we found that high TG may be a causal risk factor for depression and related traits. The results for cholesterols were more complex and mixed, but there is moderate evidence to suggest that HDL-c may be causally associated with depression traits. The findings may help shed light on the mechanisms underlying depression, and may have clinical implications. However, further clinical studies are required to replicate the findings and to investigate the effects/side-effects of lipid-lowering therapies. Further experimental works are also warranted to elucidate the mechanism involved.

## Supporting information

List of Supplementary Tables

## Acknowledgements

We would like to thank Prof. Stephen Tsui and the Hong Kong Bioinformatics Center for computing support. This study was partially supported by the Lo Kwee Seong Biomedical Research Fund, the Health and Medical Research Fund (06170506) and a Chinese University of Hong Kong Direct Grant.

## Author contributions

Conceived and designed the study: HCS. Study supervision: HCS. Main data analysis: HCS, CKLC, with advice from PCS. Data interpretation: HCS, YYC, PCS. Drafted the manuscript: HCS, YYC.

