## Supplementary material for "Causal relationships between blood lipids and depression phenotypes: A Mendelian randomization analysis": List of Supplementary Tables

Supplementary Tables are available at the journal’s website and at <https://doi.org/10.6084/m9.figshare.11574900>

Table S1 Full MR results when lipids are considered as exposure

Table S2 Results of multivariable MR (four tabs are shown; multivariable IVW and Egger in the first 2 tabs and the results at r2=0.01 shown in the last 2 tabs)

Table S3 Results of LD score regression (for genetic correlations)

Table S4 Steiger Directionality test results

Table S5 MR results of ‘negative control’ experiment in which mean platelet volume (MPV) was treated as exposure (two tabs are shown: the first shows the results at the best r2 clumping threshold; the second shows the full results)

Table S6 Full results of MR analysis in which MDD is considered as exposure

(the results for DS and DSH/suicide were the same regardless of the r2 threshold and only MR-IVW is performed due to small number of SNPs; therefore only the full results for MDD are presented)
