## Supplementary Text for "Causal relationships between blood lipids and depression phenotypes: A Mendelian randomization analysis"

### A more detailed description of GWAS study samples

### *Lipid traits*

### The study samples were primarily of European ancestry. Four lipid traits are studied, including LDL-c, HDL-c, triglyceride (TG) and total cholesterol (TC). GWAS was performed by the Global Lipids Genetics Consortium, with a total sample size (*N*) of 188,577. We downloaded summary statistics of joint GWAS analysis from <http://csg.sph.umich.edu/willer/public/lipids2013/>. Briefly, this is a meta-analysis of multiple samples of primarily European ancestry. In general, subjects known to be on lipid-lowering medications were excluded, and age, sex and PCA (reflecting ancestries) were adjusted for, although each constituent study may employ slightly different analytic approaches from the others. The details of each constituent sample, such as demographics, analytic approach, gender proportions etc. were presented in Supplementary Table 1 of ref^1^. A small proportion of subjects were non-European in ancestry (see Supplementary Table 1 of ref^1^) . The number of genome-wide significant SNPs for LDL-c, HDL-c, TG and TC were 3078, 3524, 3249 and 4169 respectively. They were subject to further LD-clumping and matching with SNPs from depression traits GWAS samples prior to MR analysis.

### For further details please refer to Willer et al.^1^.

### *Depression and related phenotypes*

We included five depression-related phenotypes as follows:

(1) Major depression disorder (MDD): We employed the results from the latest GWAS from Wray et al.^2^ Due to privacy concerns, full summary statistics are only available for a subset of subjects excluding 23andMe participants (59851 cases and 113154 controls). Please refer to Supplementary Tables 1-3 (and references therein) of Wray et al.^2^ for details of each constituent study sample. We employed this set of summary statistics for MR analysis with lipid traits as exposure. The UKBB sample (14260 of 59851 cases) included some cases from self-reporting, while others were defined by clinical records. (The exact proportion, however, was not clearly stated in the original publication^2^.)

We also performed MR with MDD as the exposure, which only requires access to the genome-wide significant SNPs; we used the ‘top 10k SNPs’ dataset derived from the entire sample (135458 cases and 344901 controls) for this analysis. Cases from the 23andMe sample were entirely based on self-reporting.

(2) Depressive symptoms (DS): GWAS results were taken from Okbay et al.^3^. This is a meta-analysis that included the MDD-PGC study (*N*=18,759) and a case-control sample from the Genetic Epidemiology Research on Aging (GERA) Cohort (*N*=56,368); it also comprised an UK BioBank (UKBB) sample made up of general population (*N*=105,739), which represented the largest proportion of the total sample (~ 58% of total sample size). Depressive symptoms were measured by a self-reported questionnaire. Age, sex and principal components reflecting ancestry were included as covariates (please refer to supplementary table 11 of ref^3^).

GWAS of phenotypes (3) to (5) was based on the UKBB sample. We downloaded GWAS summary statistics from the Neale Lab (https://sites.google.com/broadinstitute.org/ukbbgwasresults/). GWAS analysis was performed using linear models with adjustment for population stratification; details of the analytic approach is given in <https://github.com/Nealelab/UK_Biobank_GWAS/tree/master/imputed-v2-gwas> and <http://www.nealelab.is/blog/2017/9/11/details-and-considerations-of-the-uk-biobank-gwas>. For binary outcome (phenotype 5), we converted the regression coefficients obtained from the linear model to those under a logistic model, based on methodology presented in ^4^. The SE under a logistic model was derived by the delta method (see supplementary text of ref^4^ https://www.ncbi.nlm.nih.gov/pmc/articles/PMC5887138/bin/1397FileS1.pdf, equations 34 to 37). Sex and the top 10 principal components were included as covariates.

(3) Longest period of depressed/low mood: This item was based on response to a question from the self-reported questionnaire for UKBB participants. The question was ‘How many weeks was the longest period when you were feeling depressed or down?’ (data field 4609; *N* = 104,190). Response to this question was only collected from participants who indicated they have felt depressed or down for at least one whole week. Inverse-rank normal transformation was performed prior to analysis.
(4) Number of episodes with depressed/low mood: This item was based on response to the question ‘How many periods have you had when you were feeling depressed or down for at least a whole week? (*N* = 104,190) in the UKBB questionnaire (data field 4620). Again information was only collected for those who indicated having felt depressed or down for one whole week.
(5) History of deliberate self-harm (DSH) or suicide: The analyses was based on self-reported history of deliberate self-harm or suicide attempt among UKBB participants. There were 224 positive responses among 381,462 participants (according to <https://biobank.ctsu.ox.ac.uk/crystal/field.cgi?id=20002>). The low number of positive cases may be due to under-reporting of such events. The control subjects are likely mixed with positive cases, and may be considered analogous to being only weakly ‘screened’ or almost ‘unscreened’. This does not render the analysis invalid, and the use of unscreened subjects is quite common in GWAS^5^. However the power of the study will be improved if reporting bias can be eliminated.

There is no overlap between the GWAS samples for lipids and depression phenotypes, which is also supported by the lack of significant intercept from genetic correlation analysis (Table S?).

**Accounting for SNP correlations in Mendelian randomization (MR) analysis**

As described above, MR is an analytic approach to causal inference, using genetic variants as ‘instruments’ to represent the exposure. In this study, we employed the two-sample MR approach, in which the instrument-exposure and instrument-outcome associations were estimated in different samples.

We conducted MR with several different methods, including the ‘inverse-variance weighted’ (MR-IVW)^6^, Egger regression (MR-Egger)^7^ and Generalized Summary-data-based Mendelian Randomization (GSMR)^8^ approaches. We present below briefly how MR-IVW/MR-Egger takes into account SNP correlations in the analysis.

*MR-IVW and MR-Egger accounting for SNP correlations*

The IVW framework is very widely used in MR studies. Here we used an IVW approach that is able to account for SNP correlations as described in Burgess et al.^9^. Briefly, assume
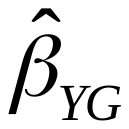
 to be the vector of estimated regression coefficients when the outcome is regressed on genetic instruments and
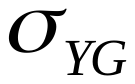
 to be the corresponding standard errors (SE), and
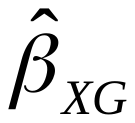
 to be the estimated coefficients when the risk factor is regressed on the genetic instruments with SE
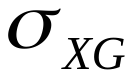
. We also assume the correlation between two genetic variants G_1_ and G_2_ to be
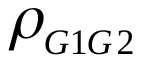
, and
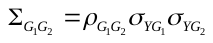
.

The estimate from a weighted generalized linear regression can be formulated by

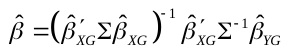

with SE

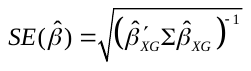

A similar approach may be used for MR-Egger, which allows an intercept term in the weighted regression. Please also refer to ref. ^7^^,^^10^ for details. The presence of imbalanced horizontal pleiotropy could be assessed by whether the intercept term is significantly different from zero.
